# EquCab3, an Updated Reference Genome for the Domestic Horse

**DOI:** 10.1101/306928

**Authors:** Theodore S. Kalbfleisch, Edward S. Rice, Michael S. DePriest, Brian P. Walenz, Matthew S. Hestand, Joris R. Vermeesch, Brendan L. O’Connell, Ian T. Fiddes, Alisa O. Vershinina, Jessica L. Petersen, Carrie J. Finno, Rebecca R. Bellone, Molly E. McCue, Samantha A. Brooks, Ernest Bailey, Ludovic Orlando, Richard E. Green, Donald C. Miller, Douglas F. Antczak, James N. MacLeod

## Abstract

EquCab2, a high-quality reference genome for the domestic horse, was released in 2007. Since then, it has served as the foundation for nearly all genomic work done in equids. Recent advances in genomic sequencing technology and computational assembly methods have allowed scientists to improve reference assemblies of large animal and plant genomes in terms of contiguity and composition. In 2014, the equine genomics research community began a project to improve the reference sequence for the horse, building upon the solid foundation of EquCab2 and incorporating new short-read data, long-read data, and proximity ligation data. The result, EquCab3, is presented here. The count of non-N bases in the incorporated chromosomes is improved from 2.33Gb in EquCab2 to 2.41Gb from EquCab3. Contiguity has also been improved nearly 40-fold with a contig N50 of 4.5Mb and scaffold contiguity enhanced to where all but one of the 32 chromosomes is comprised of a single scaffold.

---

The domestic horse *Equus caballus* is a culturally, economically, and historically important domesticated animal. Since horses were domesticated ~5kya in central Asia^1^, humans have used them extensively for agriculture, transportation, military conflict, and sport. Horses have been selectively bred for speed, strength, endurance, size, appearance traits, and temperament.

EquCab2, a high-quality reference genome assembly of the domestic horse, was released in 2007^2^. This assembly was generated using the best genomic sequencing and assembly technologies available at the time, namely: Sanger sequencing, bacterial artificial chromosome (BAC) end pairs, radiation hybrid (RH) mapping, and fluorescence in situ hybridization (FISH) mapping. Since then, many researchers have used this reference genome to study the genetics of various traits in horses^3–9^, as well as their health^10–13^ and evolution^14–17^. However, EquCab2 contains numerous gaps in scaffolds as well as sequences unassigned to chromosomes, and genomic DNA resequencing^18^ and gene annotation^19^ studies have found inconsistencies in this genome. Therefore, new genomic technologies present an opportunity to improve the equine reference genome.

We present here a new reference assembly for the domestic horse, EquCab3. This assembly benefited from rapidly evolving high-throughput sequencing technologies and new algorithms used to assemble data from these platforms. Specifically, this project began from the solid foundation of 6.8-fold coverage Sanger sequence data^2^, as well as a radiation-hybrid map and FISH data^20^. These data were augmented with 45-fold coverage Illumina short-read data that improved the characterization and accuracy of unique regions of the genome, increasing the contig N50 values 10-fold. Two different proximity ligation library preparation protocols made it possible to order these contigs and generate chromosome length scaffolds. In EquCab3, only chr6 is comprised of more than one scaffold. Finally ~16X PacBio long reads made it possible to close many of the gaps between the ordered contigs, thereby improving the contig N50 values 4-fold again. The resulting assembly is enhanced not only in contiguity but also in composition. This new version of reference sequence for the domestic horse reduces the number of gaps 10-fold and increases the number of assembled bases by 3% in the incorporated chromosomes over EquCab2.

## Results

### A new reference assembly of the domestic horse genome

We generated a new reference assembly of the domestic horse using first-, second-, and third-generation sequencing data as well as physical chromosome maps. This new reference is derived from the same female Thoroughbred horse, Twilight, that produced EquCab2. We did not attempt to derive a new mitochondrial genome sequence, and instead relied on the work done by Xu and Arnason^21^.

We used both previously published data and newly generated data to generate this reference assembly. The previously published data sets are comprised of the data used to construct EquCab2: Sanger sequencing data, BAC-end pairs^2^, and a physical map containing radiation hybrid and FISH markers^20^. For this assembly, we generated shotgun Illumina short reads, Chicago and Hi-C proximity ligation libraries, PacBio long reads, and 10x Chromium linked reads. As there is no existing software or method for creating an assembly from this combination of data types, we developed a custom pipeline to leverage the strengths of each of these data sets.

First, we used the high coverage (45x) and accuracy of Illumina short reads to generate “super-reads” with MaSuRCA^22^. We assembled these super-reads together with the long and accurate but lower coverage (6.8x) Sanger reads to create an initial assembly with Celera Assembler^23^. We scaffolded this initial assembly with the long insert-size Chicago and Hi-C proximity ligation libraries using the HiRise scaffolder^24^. To identify and correct misassemblies, we mapped all physical markers and sequence data, including BAC-end sequences, to the resulting scaffolds. We filled gaps in the corrected scaffolds with PacBio reads, which are longer but lower-accuracy than Sanger reads, using PBJelly. We phased the genome using 10x Chromium linked reads and the longranger pipeline. We aligned the high-identity and coverage Illumina short reads to the genome and used these alignments to correct errors. Finally, we used the physical map to assign scaffolds to chromosomes. The resulting assembly, EquCab3, is an improvement over EquCab2 in terms of contiguity, completeness, read mappability, and agreement with the physical map.

### Improved Contiguity

EquCab3 has improved N50 values for both contigs and scaffolds over those reported for EquCab2. For the contigs, an N50 value of 4.5 Mb vs 112 kb, and for scaffolds, 86 Mb vs 46 Mb (Table 1). At each phase of the assembly process (described in Methods), there is an improvement in either the contig or scaffold N50 over the values achieved in EquCab2. The one exception is the scaffold N50 of the Sanger + MaSuRCa Super Reads. Our scaffold N50 is 6.6Mb, less than the final value of 46Mb reported in Wade et al. (2009). The EquCab2 value incorporated additional long range data such as BAC-end reads from a library derived from Twilight’s half-brother Bravo, as well as radiation hybrid map data. With all PacBio and proximity ligation data from Twilight included, the contig N50 is increased 40-fold, and the scaffold N50 is increased from a chromosome arm-limited 46Mb to a chromosome length-limited 86Mb. Further, the total count of gaps in the ordered chromosomes is decreased more than 90%, from 42,304 in EquCab2 to 3,771 in EquCab3.

**Table 1.**

| EquCab3 Sequence Composition | Contig N50 | Scaffold N50 |
| --- | --- | --- |
| Sanger + MaSuRCa Super Reads | 1.2Mb | 6.6Mb |
| Sanger + MaSuRCa Super Reads + Chicago + HiC | 1.2Mb | 86Mb |
| Sanger + MaSuRCa Super Reads + Chicago + HiC + PacBio | 4.5Mb | 86Mb |
| EquCab2 | Contig N50 | Scaffold N50 |
| Sanger Fosmid + BAC + RH Map data | 112kb | 46Mb |

### Read mapping

The equine genome community is participating in the Functional Annotation of Animal Genomes (FAANG) project. The initial phase of this project has produced RNA-seq and whole genome shotgun sequence data from two Thoroughbred mares that are not the subject of the reference assembly. Data from both horses have been mapped to both EquCab2 and EquCab3^25^. In the first phase of the equine FAANG effort, the RNA-seq data are comprised of samples from eight tissues. As shown in Figure 1, for RNA-seq, unique mappings of the reads are increased by an average of 2.15% over EquCab2, and WGS paired reads improved by 0.44%. In the WGS dataset, more reads mapped to EquCab3 than EquCab2 (Figure 2) and the count of reads mapping in a proper pair, i.e., with both ends mapping with correct orientation, increased from a value of 811,622,501 to 814,804,213, an increase of 0.38% of the total read count.

**Figure 1:**
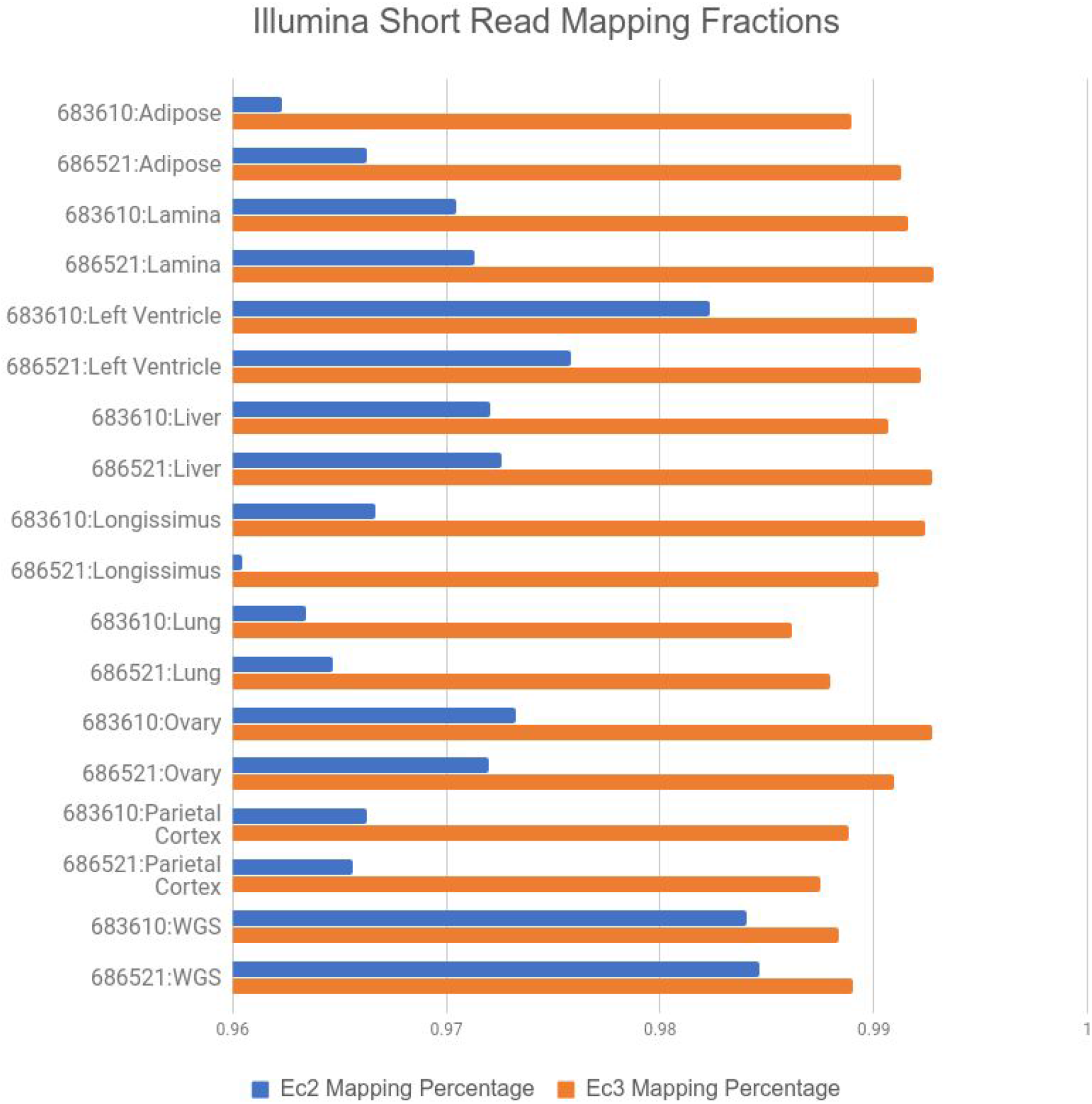
Percentages of RNA-seq reads from eight tissues from two horses and genomic reads mapping to EquCab2 vs. EquCab3. We used sequence data from FAANG for this mapping. More RNA-seq reads map to EquCab3 than to EquCab2 for every tissue in both horses. The percentage of genomic reads (last two rows; “WGS”) mapping to EquCab3 is also larger than those mapping to EquCab2, but the difference is not as large.

**Figure 2:**
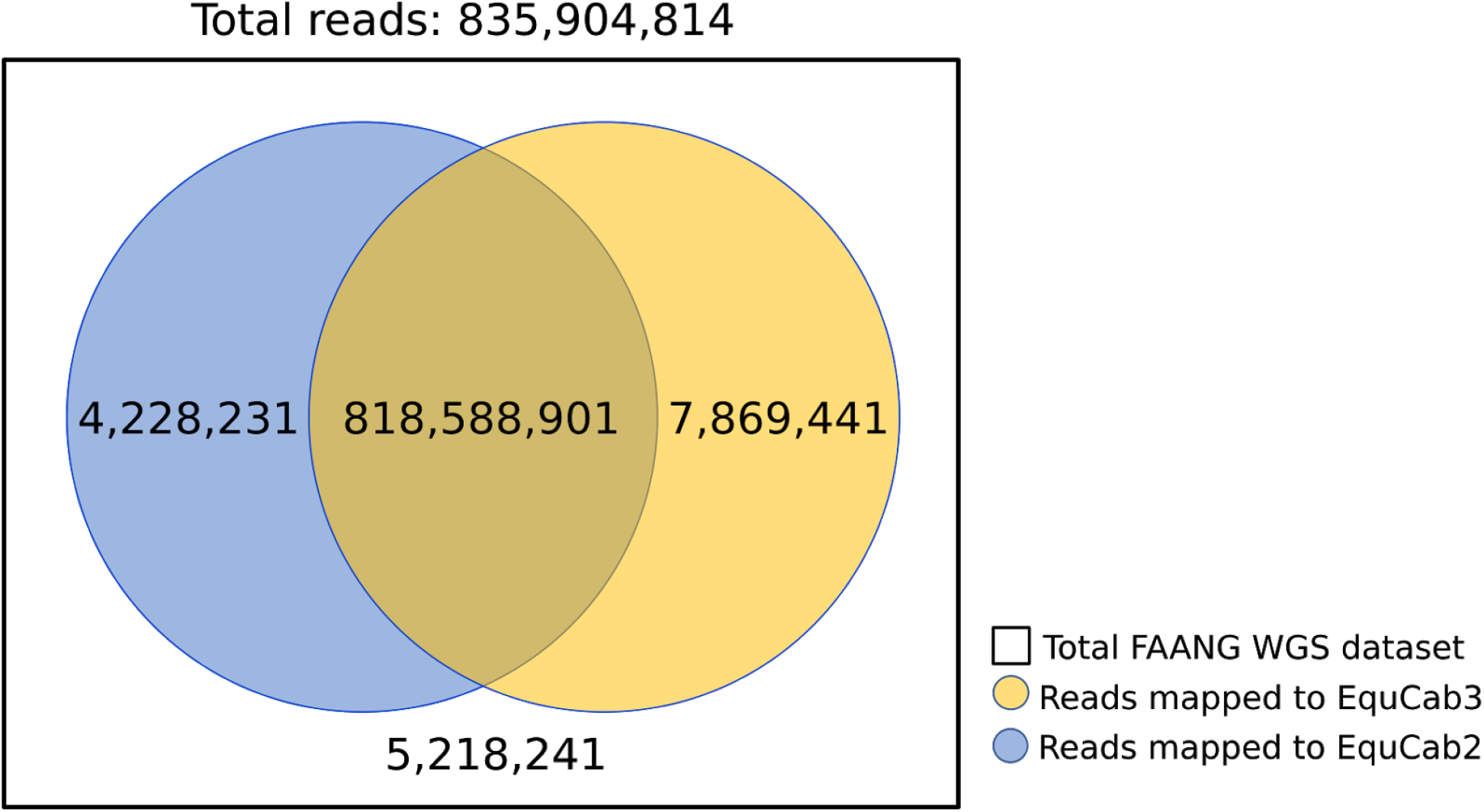
Number of reads from the FAANG WGS dataset mapping to EquCab2 and EquCab3. Significantly more reads map only to EquCab3 than only to EquCab2.

This increase in read mapping is a function of several ways in which EquCab3 is an improvement over EquCab2. EquCab3 is more accurate due to the high-coverage high-identity Illumina data used both in the initial assembly and polishing steps, and contains fewer gaps than EquCab2 due to the long read gap-filling step, resulting in fewer dips in alignment coverage, shown in Figure 3a. In addition, EquCab3 has more sequence assigned to chromosomes, giving reads more total sequence to map to, also demonstrated by Figure 3a from the length increase in chr31 from EquCab2 to EquCab3. Finally, EquCab3 improves the characterization of GC-rich regions.

**Figure 3:**
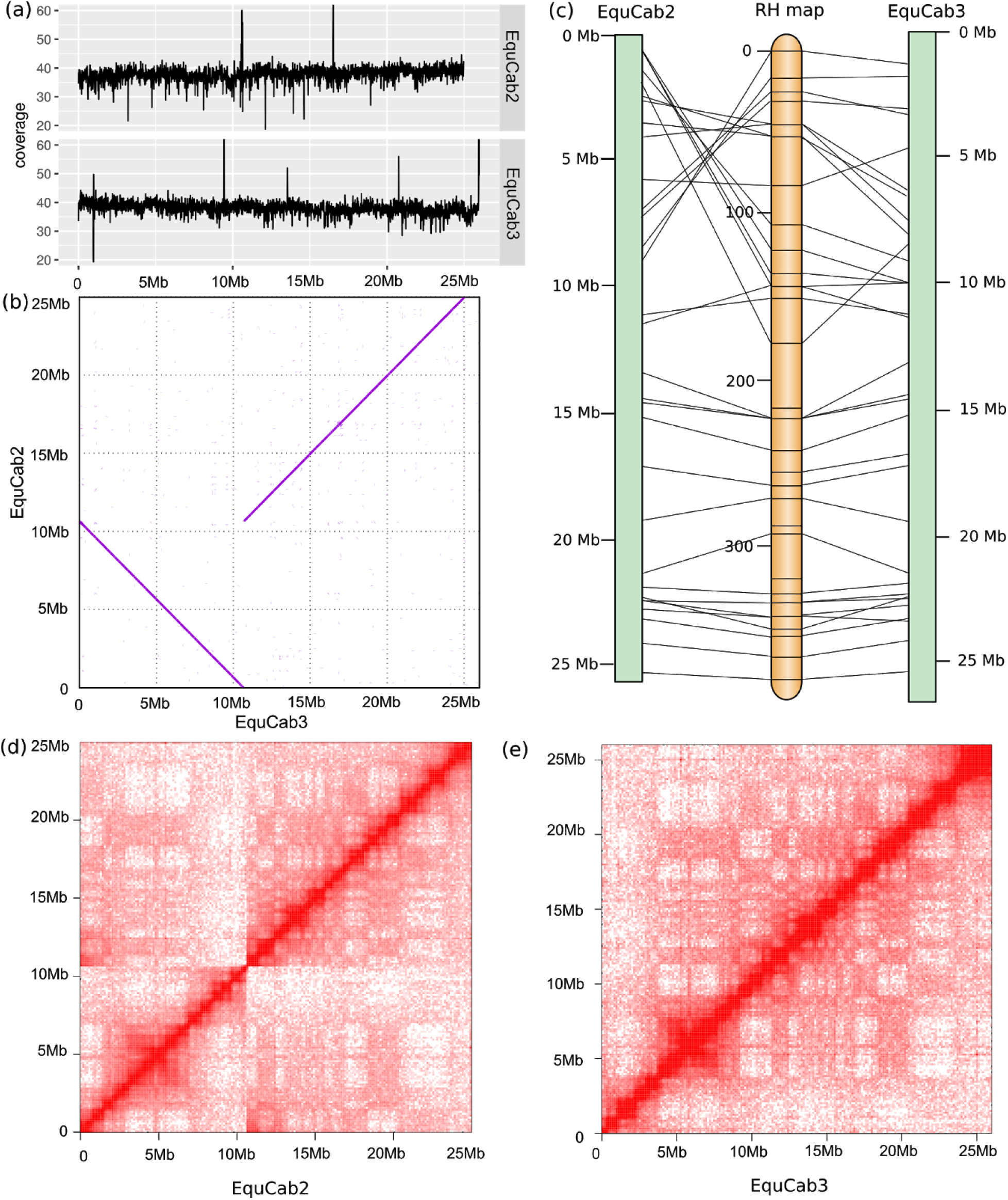
A comparison of equine chromosome 31 between EquCab2 and EquCab3. (a) Average coverage per 10kb window across chr31 in EquCab2 and EquCab3. EquCab3 has fewer coverage drops and more total sequence than EquCab2. (b) An alignment of chr31 in EquCab2 and EquCab3 shows a large inversion between the two reference genomes. The RH map (c) and Hi-C contact heat maps for EquCab2 (d) and EquCab3 (e) indicate that this discrepancy is the result of a misassembly in EquCab2.

The GC content of EquCab3 is roughly equivalent to that of EquCab2 (both near 41.6%). However, the GC fraction of the WGS reads for the two FAANG horses that mapped to EquCab3 but failed to map to EquCab2 is 48.9%. The GC content for the entire WGS dataset is 41.8%. This demonstrates an improvement in the characterization of GC-rich regions of the equine genome, and is largely attributable to the PCR-free library preparations now in common use.

We also assessed the quality of EquCab3 by aligning ancient DNA (aDNA) reads to it. EquCab2 has been used in many studies as a reference for DNA recovered from paleontological samples, giving insight into the evolution and domestication of horses^14–18^. We compared mapping statistics between EquCab3 and EquCab2 for 13 previously sequenced ancient horses^17^ (Supplementary Table S2). A paired Wilcoxon test showed a significant improvement in mapping (p=0.0017), with all 13 samples having more reads mapped to EquCab3 than to EquCab2.

### Agreement with existing RH map

We used a radiation hybrid map of the horse genome to assign scaffolds to chromosomes^20^. EquCab3 agrees with the radiation hybrid map more often than EquCab2. Of the 4,103 markers on the physical map, 2,982 map to EquCab2 while 3,039 map to EquCab3. In addition, EquCab2 contains 391 marker pairs that are oriented differently on the assembly than on the map, whereas EquCab3 contains 395, despite the 57 additional markers mapping to EquCab3. This improvement can be attributed to the lower rate of misassemblies from the use of proximity ligation data for scaffolding. An example of a misassembly in EquCab2 corrected in EquCab3 is shown in Figure 3b-e.

Of the 395 misoriented marker pairs on EquCab2, 352 are oriented the same way on both EquCab2 and EquCab3, but differently on the map. Given the multiple, orthogonal data types and differing assembly strategies used to construct EquCab2 and EquCab3, we suggest that some or all of these 352 marker pairs are oriented correctly in both assemblies but incorrectly on the RH map. Of the remaining 43 marker pairs that are misoriented on EquCab3 but not on EquCab2, 36 of these pairs do not have both markers mapping to EquCab2, leaving only 7 marker pairs agreeing with EquCab2 but not EquCab3. Given that the RH map was used to guide the assembly of EquCab2, we find this level of disagreement acceptable.

### Protein set completeness and comparative annotation

We used two methods to evaluate the completeness of our genome: universal ortholog analysis and comparative annotation. For universal ortholog analysis, we used BUSCO^26^ and the mammalian universal ortholog set. Out of 4,104 mammalian universal orthologs, BUSCO found 4,092 (99.7%) as complete orthologs in EquCab3 with 5 fragmented and 7 missing, compared to 4,064 (99.0%) complete orthologs in EquCab2 with 27 fragmented and 13 missing. EquCab3’s higher BUSCO score indicates that it is more complete than EquCab2.

Comparative Annotation Toolkit (CAT) is a software pipeline that leverages whole genome alignments, existing annotations, and comparative gene prediction tools to simultaneously annotate multiple genomes, defining orthologous relationships and discovering gene family expansion and contraction^27^. CAT also diagnoses assembly quality by investigating the rate of gene model-breaking indels seen in transcript projections from a reference as well as looking at the rate of transcript projections that map in a disjoint fashion. We performed comparative annotation of EquCab2 and EquCab3 using the genomes of pig, cow, white rhinoceros, elephant, and human. Comparative annotation of EquCab3 and EquCab2 found that more orthologs of genes in the other genomes were found in EquCab3 than in EquCab2 (Figure 4a), fewer predicted genes were split between contigs in EquCab3 than in EquCab2 (Figure 4b), and the distribution of gene coverage is significantly better in EquCab3 than in EquCab2 (Figure 4c). These results indicate that EquCab3 is a more complete and contiguous assembly than EquCab2.

**Figure 4:**
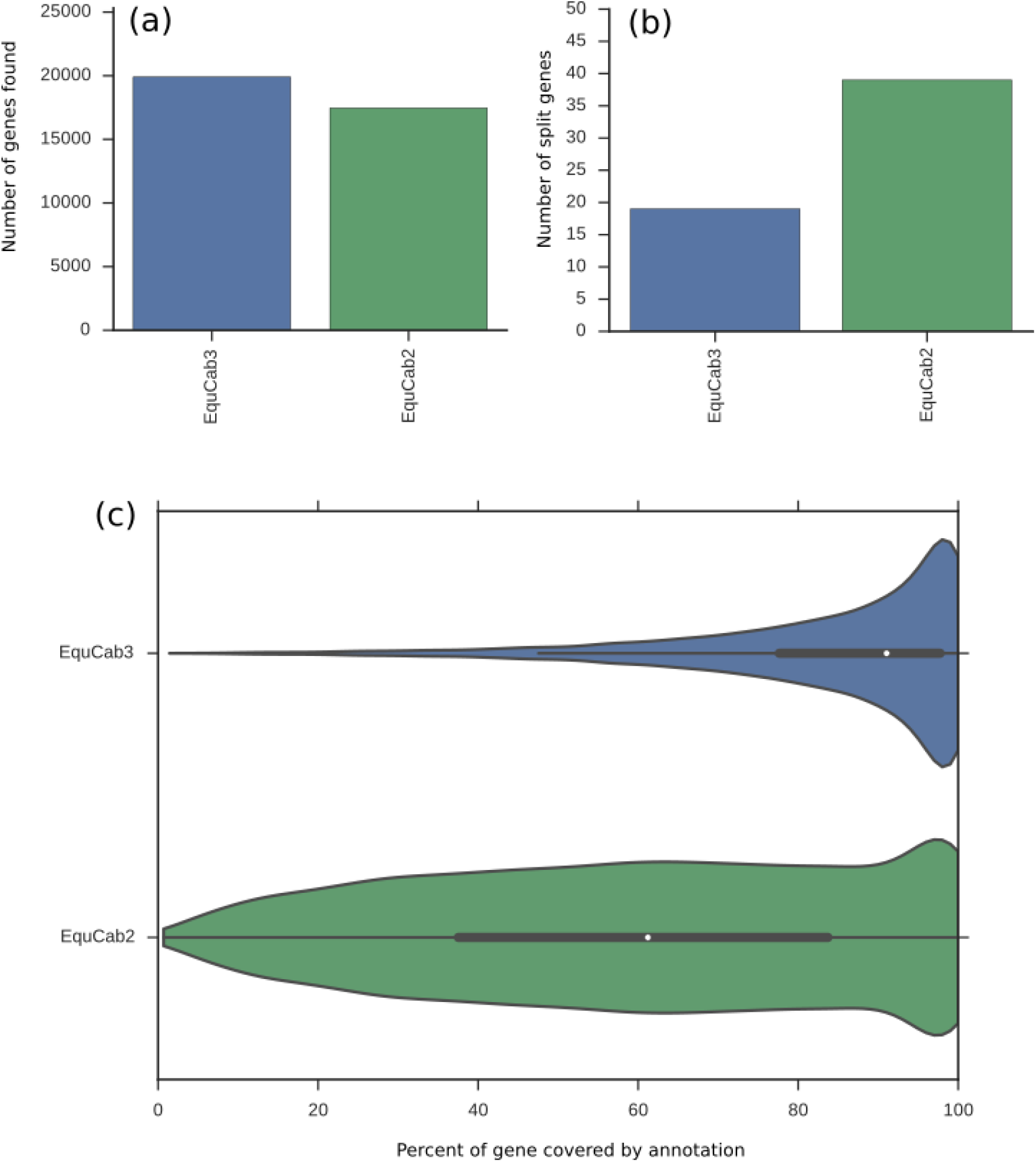
Annotation of EquCab2 and EquCab3 with the Comparative Annotation Toolkit shows substantial improvement in EquCab3. (a) More genes found in related species were annotated in EquCab3 than in EquCab2. (b) Fewer genes were split between contigs in EquCab3 than in EquCab2. (c) The gene coverage distribution is significantly better in EquCab3 than in EquCab2.

### Phasing

Most published assemblies of diploid organisms are pseudo-haploidizations produced by arbitrarily choosing between the two alleles at each heterozygous site in the genome. The 10X Chromium platform is useful for haplotype phasing, as each set of linked reads it produces comes from the same haplotype. We took advantage of this by using 10X reads and the longranger pipeline to phase Twilight’s variants in EquCab3. For each phase block inferred by longranger, rather than arbitrarily choosing which haplotype to include in the final assembly, we chose the allele which is most common among 4 Thoroughbreds, the 2 FAANG horses, and data from two other Thoroughbreds from an earlier study by Sarkar et al.^12^ This makes the reference pseudo-haploidization more similar to the population average and thus more likely to contain the ancestral allele at each heterozygous site in Twilight’s genome. For analyses which would be adversely affected by this ancestral reference bias, we provide the phased 10X variant calls as supplemental data.

## Discussion

This new genome represents an improvement for the horse reference in terms of both composition and contiguity. It is also more consistent with the existing RH map and FISH data for the horse than was EquCab2. Going forward, the lens through which this reference will be viewed will be as an alignment target for the vast amount of high throughput sequence data that will continue to be generated for the horse and other related species. The assembly process described here was guided and informed by data that included not only high quality short reads but long reads and proximity ligation data. All equine data produced by any of these technologies should be well served going forward. The most common data types for genetic and genomic studies, Illumina short reads, have been demonstrated to map to the new reference for two non-related Thoroughbreds at an improved average rate of 2.15% for RNA-Seq and 0.44% for WGS libraries. In a comparative genomics analysis, more gene orthologs were found, and for those that were found, the coverage of the homologous transcript sequence was more complete. The new long-range sequence data not only improved the contiguity of the genome, but allowed us to phase the genomic data for Twilight. Finally, the regions added for the genome were higher in GC content, which will enable a better characterization of both genetic variation and epigenetic status in GC-rich regulatory regions for the horse.

This represents a culmination of a project conceived and begun in 2014 with the support of the equine genomics community. Although it will certainly not be the last reference genome for the domestic horse produced for public annotation, it should foster genetic and genomic discoveries for years to come.

## Methods

### Sequence data Generation

**Sanger Data:** The Sanger sequence data for the Thoroughbred mare Twilight, produced for and used to build EquCab2^2^, were downloaded from the NCBI Trace Archive as described in Rebolledo-Mendez et al.^18^

**Illumina PE HiSeq and MiSeq:** Construction of a PCR-free shotgun genomic library and sequencing on MiSeq and HiSeq2500 instruments were carried out at the Roy J. Carver Biotechnology Center, University of Illinois at Urbana-Champaign (UIUC).

A shotgun genomic DNA library with an insert size of 500bp (range 300 - 650bp) was constructed from 2μg of Twilight’s genomic DNA after sonication with a Covaris M220 (Covaris, MA) with the Hyper Library Preparation Kit from Kapa Biosystems (Roche) with no PCR amplification. The adaptored DNA library was loaded onto a 2% agarose gel and fragments 450bp to 550bp in length were cut from the gel and recovered with the QIAquick gel extraction kit (Qiagen, CA). The size-selected library was quantitated with Qubit (ThermoFisher) and run on an Agilent bioanalyzer DNA high-sensitivity chip (Agilent, Santa Clara, CA) to confirm the presence of DNA fragments of the expected size range. It was further quantitated by qPCR on a BioRad CFX Connect Real-Time System (Bio-Rad Laboratories, Inc. CA) prior to sequencing for maximization of the number of clusters in the sequencing flowcell.

The PCR-free shotgun library was first sequenced on a MiSeq with v3 reagents to generate paired-reads 300nt in length. The data confirmed the DNA fragment sizes. The library was subsequently sequenced on a HiSeq2500 for 161 cycles from each end using a TruSeq Rapid SBS sequencing kit1 v1. The fastq read files were generated with the bcl2fastq v1.8.4 Conversion Software (Illumina, San Diego, CA).

**PacBio:** Ten micrograms of high molecular genomic DNA from Twilight was sheared with gTUBES (Covaris) in an Eppendorf^®^ 5424 centrifuge at 4800 RPM for 2x 60 seconds. A single PacBio library was prepared from this following PacBio’s protocol P/N 100-286-000-07 (20 kb Template Preparation Using BluePippin(Tm) Size-Selection System) with PacBio DNA Template Prep Kit 1.0. For the size selection, the sample was run on a 0.75% BluePippin cassette (ref: PAC20KB) using the pre-defined ‘0.75% DF Marker S1 high-pass 6-10kb vs3’ program and a cut-off of 10-50kb. The library was sequenced on 88 SMRT cells on a PacBio RSII using

DNA/Polymerase Binding Kit P6 and DNA Sequencing Kit 4.0 (v2) sequencing reagents, magbead loading, and stage start. All SMRTcells were run through PacBio’s SMRT Portal v2.3.0 pipeline RS_subreads.1 with default settings except for minimum subread and polymerase read lengths of 1kb. In addition, reads-of-insert were generated using the RS_ReadsOfInsert.1 pipeline with a minimum insert read length set to 1kb. Reads-of-insert had a mean of 4 passes and length of 11,785 bp.

Of the total initial read count, 5,934,426, we were able to create circular consensus (.ccs) reads totalling 371,943 reads. The remainder of the reads were used to generate a reads of insert file consisting of 5,562,483. These two datasets were used in the PBJelly runs described below.

**CHiCago library:** We generated a CHiCago library as previously described^24^ using blood from Twilight.

**Hi-C library:** We generated a Hi-C library with primary fibroblasts from Twilight using a Hi-C protocol modified such that the chromatin immobilization took place on magnetic beads. We crosslinked the fibroblasts in formaldehyde, and lysed, washed, and resuspended as described by Lieberman-Aiden et al.^28^ We then immobilized the chromatin on SPRI beads as described by Shendure et al.^29^ We restriction digested the DNA with DpnII, labeled ends with biotinylated dCTP, ligated ends, and reversed crosslinks. The sample was prepared for sequencing using the NEB Ultra library preparation kit according to the manufacturer’s instructions, with one exception: prior to indexing PCR, the sample was enriched by pulldown on 30 μL Invitrogen C1 Streptavidin beads, then washed to remove non-biotinylated DNA fragments.

**10X Genomics library:** Twilight’s genomic DNA was size selected for fragments >40 Kbp on a BluePippin instrument (Sage Sciences, Beverly MA) and Illumina sequencing libraries were constructed using the 10X Genomics Chromium Controller instrument with their Genome Reagents Kit v2 chemistry (10x Genomics, Pleasanton CA) according to the manufacturer’s recommendations. The resulting Illumina library was sequenced on a NextSeq500 using a High Output Kit v2 for a paired-end, 2x151 bp run (Illumina, San Diego CA). The data were analyzed and assembled using the 10x Genomics Supernova version 1.1.5 pipelines.

### Assembly Generation

**MaSuRCA:** The 42X Illumina PE data described above was assembled into super-reads using MaSuRCA^22^ version 3.1.3. The super reads produced a reduced representation of fragments with ~2X coverage (4.7Gb) with a contig N50 of 1,734 nucleotides.

**Celera Assembler:** The Celera Assembler^30,31^, version 8.2 (downloaded from http://wgs-assembler.sourceforge.net/wiki/index.php?title=Main_Page) was used to create contigs and scaffolds using the super reads produced by MaSuRCA and the EquCab2 Sanger sequence data.

**HiRise:** We scaffolded the output of Celera Assembler using HiRise version 2.1.1 in serial mode with default parameters, with the CHiCago and Hi-C libraries as input libraries^24^.

**Identifying misassemblies:** In order to identify misassemblies in the HiRise assembly relative to EquCab2, we aligned the HiRise output scaffolds to EquCab2 using nucmer with default parameters^32^. In every place where the alignment indicated a difference in order and orientation of scaffolds between the two assemblies, we used every available data type to resolve the discrepancy and determine which was correct. Our strategies included aligning BAC-end pairs from a half-brother of Twilight^2^ to the assemblies using bwa mem with default parameters^33^, assessing concordance with the physical map, looking for split genes predicted by the Comparative Annotation Toolkit^27^, aligning coding sequences of any genes in the region to the assemblies using gmap with default parameters^34^, and examining heatmaps of long-range read pairs mapping to the assembly generated by the HiRise and longranger pipelines.

**PBJelly:** We filled gaps in the manually corrected HiRise scaffolds from the previous step using PacBio reads of insert and circular consensus reads as input to the PBJelly (version PBSuite_15.8.24) pipeline with the steps setup, mapping, support, extraction, assembly, and output, in that order.

**Assigning scaffolds to chromosomes:** We used a previously published radiation hybrid map^20^ to assign scaffolds to chromosomes. We aligned each physical marker’s STS primers to the assembly using bwa fastmap^35^ and used only markers with both primers aligning uniquely and in the correct orientation. We then placed scaffolds on chromosomes based on the markers’ mapping locations.

### Quality control and assessment

**Read Mapping:** Short read sequence data generated in the initial phase of the equine Functional Annotation of Animal Genomes (FAANG) project has been mapped to both EquCab2 and EquCab3 for comparison of mapping fractions. Both Whole Genome Shotgun (WGS) sequence (40X), and RNA-Seq (avg 20M reads/tissue) datasets from 8 tissues types for each of two animals were trimmed using TrimGalore (a wrapper for Cutadapt^36^). For WGS data, the program BWA^35^ (version 0.6.1) aln module was used to align the reads to the reference. BWA sampe was used to produce a usable SAM file. SAMtools^37^ (version 0.1.18) was used to convert from SAM to BAM format. Picard (version 1.65) FixMateInformation and MarkDuplicates modules were used, followed by GATK^38^ (version 1.5) RealignerTargetCreator, and IndelRealigner (validation_strictness set to LENIENT for each). For the RNA-seq data, the mapping program STAR^39^ (version 2.5.3a) was used with default parameters except for the following: –readFilesCommand zcat –outSAMtype BAM SortedByCoordinate –outBAMsortingThreadN 16 -outSAMunmapped Within.

**Polishing:** Since Twilight’s sequence data and the EquCab3 were derived from the same animal, any homozygous differences between the PE data and the reference of which they are a component are likely errors. The differing bases were likely contributions from the sequence data generated on other platforms used for the assembly such as the Sanger or PacBio data.

The errors are either with the reference or with the miscalled/undersampled genotypes derived by the variant discovery software. To evaluate these positions, we performed variant discovery and genotyping with the UnifiedGenotyper using the Twilight PE data, the two FAANG thoroughbreds, and two additional thoroughbreds from Sarkar et al.^12^ whose data was downloaded from the Sequence Read Archive (BioSample/experiment accession numbers SAMN03838869/SRX1097022, SAMN03838867/SRX1097495) and mapped as described above. The UnifiedGenotyper was used in discovery mode on the cohort. The resulting variant call format file was then parsed with custom java software looking for positions at which the Twilight data produced a homozygous genotype differing from the reference. The genotypes for the other animals were then queried at those positions. If the reference allele was detected in one of the other horses, the reference nucleotide at that position was not changed.

**Removal of Microbial Contamination:** To build microbial sequence databases, all bacterial, viral, and fungal reference genomes were downloaded from RefSeq. For each of the three databases (bacteria, viruses, and fungi), the sequences were first masked with DustMasker^40^. Kraken v1.0^41^ was used to generate k-mers (k=32) and to search the EquCab3 contigs for exact matches. Contigs with at least one exact 32-mer match were considered microbial contaminants and removed from the reference sequence. A total of 41 contigs were removed in this way.

**Removal of Small Contigs:** All scaffolds smaller than 3000 bases in length were removed from the assembly that was submitted for annotation. The contig and scaffold N50s for what was submitted were 4.73Mb and 87.2Mb, respectively.

**Phasing with 10X data:** The data generated for Twilight on the 10X platform described above was mapped to the reference using the longranger (version 2.1.3) wgs module. The phased variant file produced was then used to modify individual variant positions to conform to the haplotype whose allele was most common among the FAANG horses, and two other thoroughbreds described above in *Polishing*.

**N50 calculation:** The PBJelly (version PBSuite_15.8.24) utility summarizeAssembly.py was used to calculate N50 values. The default setting of 25 was used for the minimum gap setting. This ignored any gaps sized less than 25 Ns.

**Universal ortholog analysis:** For universal ortholog analysis, we used BUSCO^26^ version 3.0.2 in protein mode with the lineage dataset mammalia_odb9 version 2016-02-13. For protein set inputs, we used the official NCBI protein sets for EquCab2.0 (accession GCF_000002305.2) and EquCab3.0 (accession GCF_002863925.1).

**Comparative annotation:** For this analysis, a progressiveCactus^42^ alignment of equCab2 and equCab3 was performed with pig (susScr3), cattle (bosTau8), white rhinoceros (cerSim1), elephant (loxAfr3) and human (hg38). The guide tree was (((Human:0.164501,((Pig:0.12,Cow:0.16908)1:0.02,(equCab3:0.0001,equCab2:0.0001):0.059397, White_rhinoceros:0.05)1:0.060727)1:0.032898)1:0.023664,Elephant:0.155646);, putting EquCab2 and EquCab3 under the same node with a branch length of 0.0001. CAT^27^ was then run using the Ensembl V89 annotation of pig as the source transcript set. No RNA-seq data were provided, and so no transcript cleanup steps or comparative gene predictions were performed. Split gene analysis is performed by looking at transcripts which have multiple projections after paralog resolution and which have multiple projections whose start and stop points are within 10bp of each other in source transcript coordinates.

**Read Filtering and Counting:** Custom java software using the htsjdk (version 2.12.01) was written to filter the mappings that were not primary (getNotPrimaryAlignmentFlag() is false) from the mapped read (getReadUnmappedFlag() is false) count.

**Ancient DNA mapping:** We downloaded single-end Illumina reads produced by a previous study^17^ (Supplementary Table S2, NCBI Bioproject PRJEB19970). Adapters and PCR artifacts were trimmed using AdapterRemoval v2^43^. For normalization across samples, fastq files were downsampled to 6M reads using seqtk (https://github.com/lh3/seqtk). Low complexity sequences were removed using PRINSEQ^44^ following bwa mapping^35^ with parameters optimized for aDNA: *aln* algorithm, “seed disable” flag, and minimum mapping phred quality of 20.

### Data Availability

The sequence read datasets generated during the current study are available in the NCBI SRA repository under accession SRP126689. The final assembly generated during the current study is available in the NCBI Genbank repository under accession GCA_002863925.1. We also provide intermediate assemblies produced during the process, a de novo assembly based solely on the PacBio data, and phased variant calls from the 10X longranger pipeline as supplementary data.

Table 2 attached as Excel Sheet

## Additional Information

### Acknowledgements

This work was supported by Morris Animal Foundation grant D15EQ-019 and the NRPS8 Horse Genome Coordinator Fund. E.S.R. is an ARCS scholar. Support for C.J.F. was provided by the National Institutes of Health (NIH) (1K01OD015134 and L40 TR001136). Alignment and assembly work for this project were performed on the University of Louisville Cardinal Research Cluster. The authors are grateful to Mr. Harrison Simrall for his assistance in installing and running many of the applications used in this work. We also wish to thank Wim Meert for PacBio runs and library preparations, and Peter A. Schweitzer who prepared the libraries for the 10X Genomics data generated for the project. Finally, we would like to thank Drs. Tomas Bergström, Sofia Mikko, Agnes Viluma, Göran Andersson, and Petr Horin for their insightful review of the MHC locus for this assembly.

### Author contributions

T.S.K., E.S.R., and B.P.W. performed the assembly. J.N.M., M.S.H., J.R.V., B.L.O., and E.S.R. contributed genomic DNA sequencing and primary data generation. T.S.K., E.S.R., M.S.D., I.T.F., and A.O.V. performed quality control on the assembly and evaluated its completeness and accuracy. J.L.P, C.J.F, and R.R.B. prepared and sequenced DNA and RNA for annotation and quality control purposes. J.N.M. designed and led the project with critical input from T.S.K., D.F.A., D.C.M., R.E.G., E.B., L.O., S.A.B., and M.E.M. T.S.K., E.S.R., M.S.D., and J.N.M. wrote the manuscript. All authors reviewed and edited the manuscript.

